# Layered Structure of Cortex Explains Reversal Dynamics in Bistable Perception

**DOI:** 10.1101/2023.09.19.558418

**Authors:** Kris Evers, Judith Peters, Rainer Goebel, Mario Senden

## Abstract

Bistable perception involves the spontaneous alternation between two exclusive interpretations of a single stimulus. Previous research has suggested that this perceptual phenomenon results from winnerless dynamics in the cortex. Indeed, winnerless dynamics can explain many key behavioral characteristics of bistable perception. However, it fails to explain an increase in alternation rate that is typically observed in response to increased stimulus drive and instead predicts a decline in alternation rate. To reconcile this discrepancy, several lines of work have augmented winnerless dynamics with additional processes such as global gain control, input suppression, and release mechanisms. These offer potential explanations at an algorithmic level. But it remains unclear which, if any, of these mechanisms are implemented in the cortex and what their biological substrates might be. We show that the answers to these questions lie within the architecture of the cortical microcircuit. Utilizing a dynamic mean field approach, we implement a laminar columnar circuit with empirically derived interlaminar connectivity. By coupling two such circuits such that they exhibit competition, we are able to produce winnerless dynamics reflective of bistable perception. Within our model, we identify two mechanisms through which the layered structure of the cortex gives rise to increased alternation rate in response to increased stimulus drive. First, deep layers act to inhibit the upper layers, thereby reducing the attractor depth and increasing the alternation rate. Second, recurrent connections between superficial and granular layers implement an input suppression mechanism which again reduces the attractor depth of the winnerless competition. These findings demonstrate the functional significance of the layered cortical architecture as they showcase perceptual implications of neuroatomical properties such as interlaminar connectivity and layer-specific activation.

**Author summary:** In our study, we explore the mechanistic underpinnings of bistable perception, a phenomenon where a single visual stimulus can be perceived in two distinct ways, and where our percept alternates spontaneously between interpretations. Although winnerless competition mechanisms have been widely recognized to govern this, they fall short in explaining why we observe more perceptual alternations with a stronger stimulus. To uncover the cortex’s role in this discrepancy, we constructed a detailed model that mirrors the layered structure and interlaminar connections of the cortex. Remarkably, the architecture of these layers emerged as instrumental players. We discovered that the deeper layers of the cortex seem to inhibit the upper layers, facilitating a quicker alternation between perceptions when stimulated. Additionally, the interlaminar recurrent connections between the upper ‘output’ layer and middle ‘input’ layer appeared to destabilize the prevailing interpretation of the stimulus, leading to faster alternations. Our research illuminates how the complex architecture of the cortex, particularly the interconnections between its layers, plays a pivotal role in influencing our perception. The layered structure of the cortex goes beyond mere anatomy; it influences our perceptual experiences.

## Introduction

Bistable perception refers to the phenomenon wherein the subjective experience of an observer spontaneously alternates between two mutually exclusive interpretations of the same physical stimulus [9, 34]. This phenomenon is integral to the study of visual perception as it provides insights into the neural mechanisms that underlie perceptual decision-making and awareness [36, 48, 54, 57]. Behaviourally, bistable perception is typically characterized in terms of dominance duration and alternation rate. Dominance duration is the duration for which one interpretation is maintained whereas alternation rate is the number of perceptual alternations within a predefined time interval.

Statistical generalities of these behavioral measures of bistable perception have been consolidated as Levelt’s four propositions [11, 35]. The first two propositions pertain to dominance duration statistics and state that 1) the interpretation receiving larger stimulus drive (stronger evidence) exhibits longer dominance durations and that 2) dominance duration increases monotonically with stimulus drive (Fig 1A). The third and fourth proposition pertain to alternation rate. Specifically, the third proposition states that alternation rate is maximal when both interpretations receive equal stimulus drive and reduces as stimulus drive diverges between interpretations [41] (Fig 1B). The fourth proposition states that when the total stimulus drive is increased but equal for both interpretations, the alternation rate increases (Fig 1B). These rules are instrumental for validating models concerning neural mechanisms underlying bistable visual perception [11, 15, 29].

**Fig 1.**
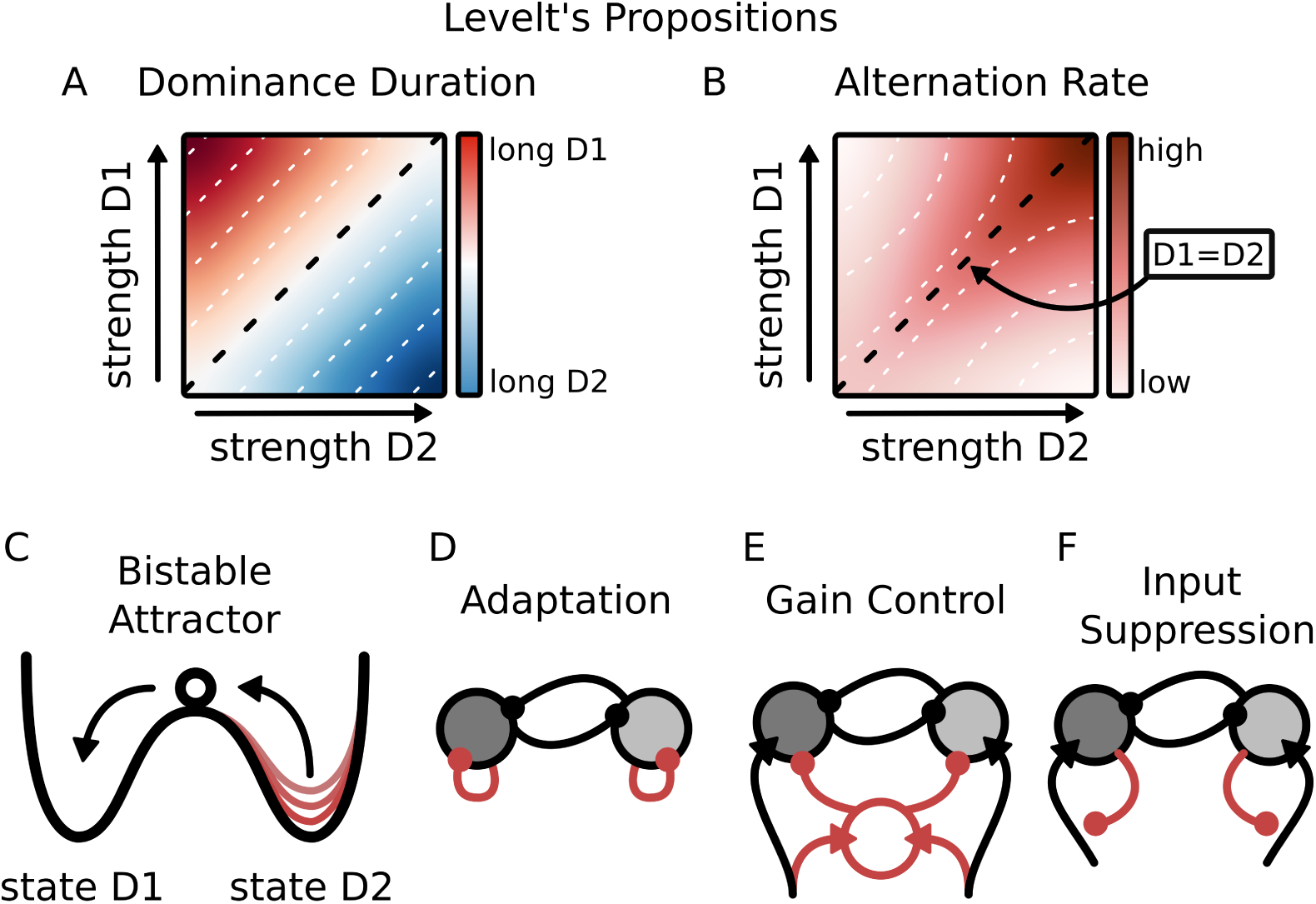
Conceptual illustration of Levelt’s propositions for behavioral statistics of bistable perception. A: Dominance duration is longer for the interpretation receiving highest stimulus drive (rule 1). Dominance duration of stronger stimulus mainly increases (rule 2). B: Alternation rate is highest when stimulus drive to both interpretations is equal (rule 3) and increases when the total drive increases (rule 4). C: double-well diagram for a bistable attractor system; The system can occupy one of two attractor states at a time (D1 or D2). Switches between states occur through noisy jumps. Depth of the attractor determines the probability of transitions; Several mechanisms can be used to maintain shallow attractors: D: Neural adaptation in populations inhibits its own activation; E: A gain control mechanism suppresses neural activity when total stimulation is increased through a shared neural component; F: Input suppression through direct suppression of the selective input.

Previous research has suggested that bistable perception results from a form of mutual inhibition between two neuronal populations reflecting the two interpretations [16, 40, 53]. Strong competition between these populations will generate winner-take-all dynamics with two attractor states corresponding to the two interpretations of the stimulus. By introducing mechanisms such as neural adaptation it is possible to achieve winnerless competition dynamics where the model switches between attractor states. Dynamics of such attractor systems are commonly conceptualized using double-well energy landscapes, where the depth of a well determines the probability of transitions from one attractor state to another [12] (Fig 1C). Winner-take-all dynamics is characterized by deep wells that render transitions highly improbable. In winnerless competition, on the other hand, attractor wells are sufficiently shallow for transitions to occur regularly, albeit stochastically. Indeed, the depth of attractor states directly determines dominance duration and alternation rate [12].

While models implementing winnerless competition have successfully replicated most of the key behavioral characteristics of bistable perception [11, 15, 40], they have faced challenges in replicating the increase in alternation rate with increased stimulus drive as posited by Levelt’s fourth proposition. A fundamental property of these models is that increasing total stimulus drive deepens attractor wells and hence *reduces* alternation rate [40, 53] (Fig 1D). In order to address this limitation, some models introduce additional mechanisms such as global gain control, input suppression, or a release mechanism. A global gain control mechanism suppresses the total neural activity when stimulation is increased by inhibiting all neuronal populations equally [40] (Fig 1E). An input suppression mechanism ensures that the currently dominant state inhibits its own input, thereby promoting transitions away from that state [15, 18, 29] (Fig 1F). A release mechanism may be utilized by the suppressed population to overcome the dominant population [16]. These mechanisms serve to maintain shallow attractors when the total input received by both populations is increased and offer explanations for Levelt’s fourth proposition at an algorithmic level [37]. However, at present it is not clear which, if any, of these mechanisms are implemented in the cortex and what might be their biological substrates.

We suggest that the answers to these questions lie within the architecture of the cortical microcircuit. The cortex is characterized by a multi-layered structure with complex interlaminar connectivity [19, 27, 49]. The horizontal organization of the cortex into columns is characterized by responses to specific sensory features [42] while the vertical organization into layers is marked by a canonical connectivity pattern that is consistent across the cortex [4, 13, 58]. As such, mutually exclusive interpretations of the same physical stimulus are represented in distinct cortical columns (c.f. [51]) that compete via layer-specific horizontal connections located primarily in superficial layers 2 and 3 (L23; [1, 3, 38, 49]).

Like several previous studies (e.g., [5, 15, 40]), we hypothesize that bistable perception is a form of winnerless dynamics, where an adaptation mechanism ensures shallow attractors and random noise causes transitions between attractor states. However, in classical winnerless competition models, the stimulus input drives directly the populations engaged in mutual inhibition. In contrast, the layered cortical circuitry in our model separates the input from the mutual inhibition mechanism. More specifically, feedforward input primarily arrives in layer 4 (L4; [6, 20, 26, 27, 47]). In contrast to L23, L4 selectively targets intra-columnar populations with similar preferred features, thus lacking lateral connectivity to surrounding cortical columns [26, 33, 62]. However, L4 projects to L23 [8, 47] providing input to winnerless dynamics between cortical columns. Finally, L23 excitatory neurons also project to L4 inhibitory neurons [14, 56]. Through these projections L23 effectively inhibits its own input. We therefore hypothesize that an input suppression mechanism is inherent in the recurrent connections between L4 and L23.

We hypothesize that the implementation of shallow attractors is further supported by a gain control mechanism implemented by the deep layers of the cortex. Activation of layer 6 (L6) in mouse cortex has been shown to strongly suppress neural activation of the upper layers [10, 21, 43]. Similarly, deep layer 5 (L5) suppresses activity in superficial and granular layers L23 and L4, respectively [44]. We suggest that L4 activates L23, next L23 activates deep layer L5, and finally deep layers L5 and L6 inhibit superficial and granular layers L23 and L4 (c.f. [47]) to adjust their sensitivity to input. Given that both L5 and L6 also receive feedforward thalamic input [2, 8, 17, 27, 28, 52], this gain control mechanism is also responsive to stimulus drive.

We test these hypotheses *in silico* using a biologically-derived layered columnar model (*Laminar Column Model*). The model comprises two columns of four layers (L23, L4, L5, L6), representing the two competing interpretations of a bistable stimulus (e.g., horizontal and vertical motion in an ambiguous motion paradigm). Each layer consists of one excitatory and one inhibitory population. Estimations of layer specific sizes of neuron populations are taken from a macaque brain atlas [50]. Moreover, we used empirically derived interlaminar connectivity [8, 47, 55]. Inspired by a recent study demonstrating that two mutually exclusive interpretations of an ambiguous motion stimulus were reflected by distinct cortical columns in (human) MT [51], we report results for interlaminar connectivity specifically of (macaque) MT. However, our results generalize to other cortical areas as well (See S1 Fig). Dynamics of each population are described by a set of dynamic mean-field equations [24]. Mutual inhibition between columns is implemented within L23 to promote competition. Noise and adaptation are included to promote state reversals between attractors (see *Laminar Column Model* diagram in Fig 2).

**Fig 2.**
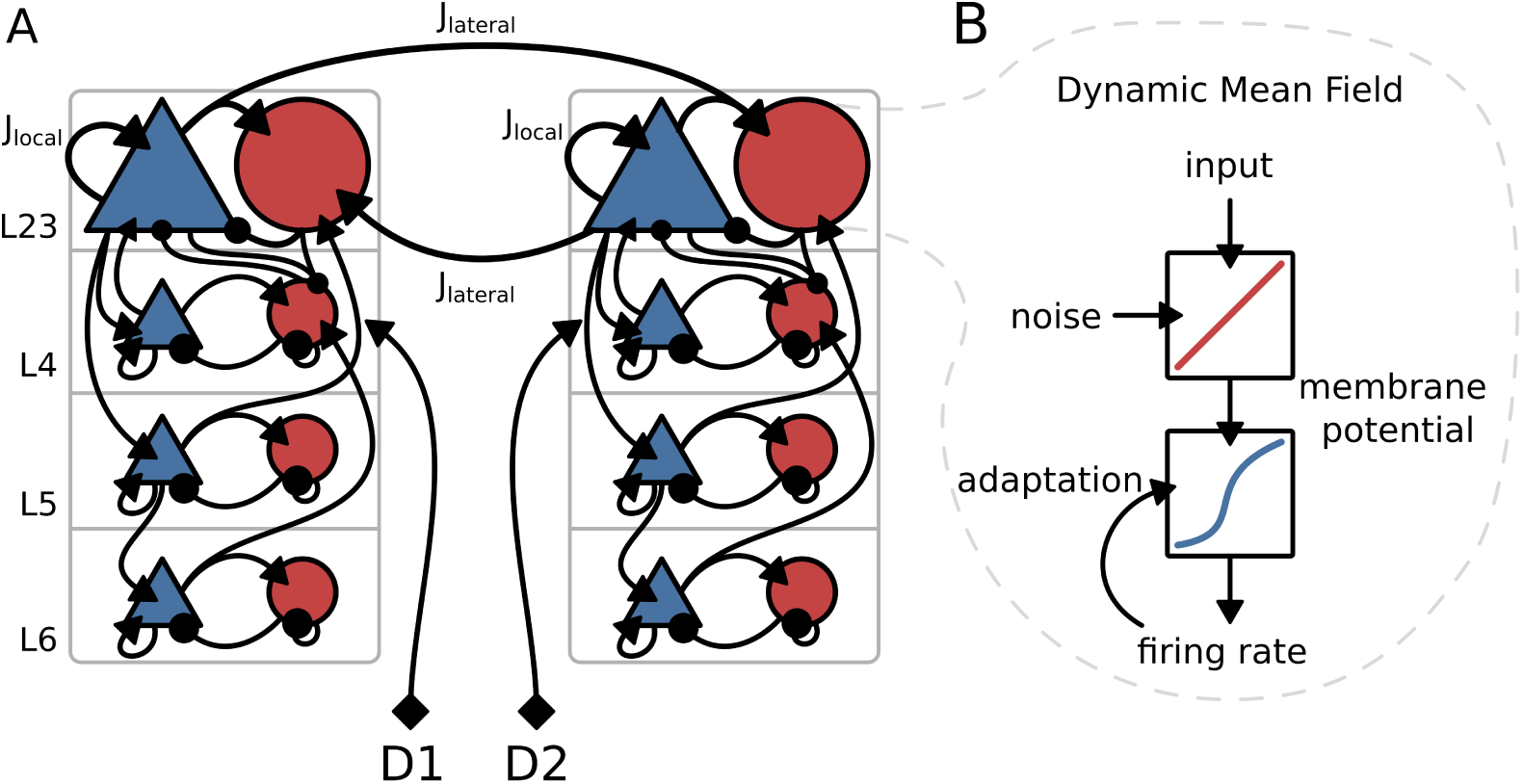
*Laminar Column Model* diagram. A: Two layered columns mutually inhibit each other through lateral connections in L23, from excitatory to inhibitory populations. *J_lateral_*and *J_local_*scale the lateral inhibition and the local self-excitation weights, respectively. L23 populations are recurrently connected to deeper layers according to the empirically derived interlaminar circuit connectivity from [47]. Each column consists of four layers (L23, L4, L5, L6). The two columns receive selective input in layer L4 reflecting the strength of the stimulus for the two interpretations (D1 & D2; e.g. horizontal or vertical apparent motion). B: The local dynamic mean field model. Each population in the network is simulated using a set of non-linear differential equations to obtain the synaptic input, the membrane potential and eventually the firing rate.

We show that the layered architecture allows for replication of all four of Levelt’s propositions. Specifically, we show that the recurrent interlaminar connectivity between superficial layer L23 and granular layer L4 implement an input suppression mechanism. This results in a suppression of L4 by L23 for strong feedforward activation of L4 and hence maintains shallow attractors. Furthermore, we show that deep layers L5 and L6 indeed implement a gain control mechanism on the upper layers. This additionally contributes to maintaining shallow attractors. Both input suppression and gain control mechanisms in the empirically derived columnar model ensure full replication of Levelt’s propositions.

These findings highlight the functional significance of the layered cortical architecture. We show that the cortex implements two of the proposed mechanisms that ensure shallow attractors in spite of strong stimulus drive. This model sets the stage for future research endeavors that aim to elucidate the functional role of laminar cortical circuits for perception [27].

## Results

### The *Laminar Column Model* reflects empirically observed Bistable Perception Dynamics

It has previously been shown that self-excitation of L23 excitatory neurons controls the dynamical stability of the columnar model [7]. We extend these findings and show that self-excitation in L23 is also important for the competitive dynamics of the column. Specifically, the dynamic range of the model is primarily determined by the strength of self-excitation within L23 of one column (*J_local_*) and the strength of lateral inhibition between L23 across columns (*J_lateral_*) Feedforward stimulation of L4 enables competition between columns (Fig 3A). Higher values of both self-excitation and lateral-inhibition promote competition between the columns (Fig 3A). To simulate winnerless competition dynamics, in the following we take parameter values within the winnerless regime that are biologically realistic (see Methods for details).

**Fig 3.**
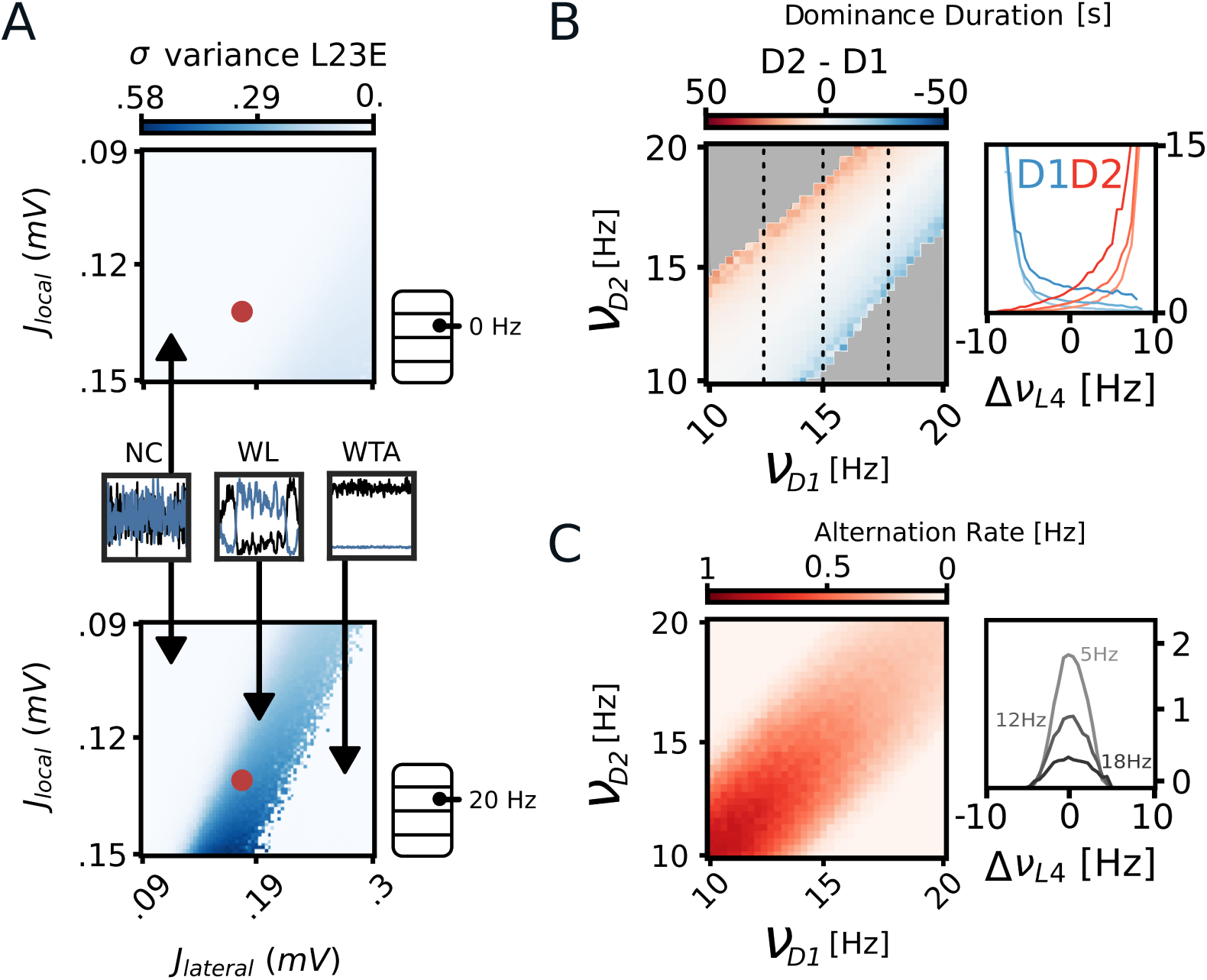
Exploring winnerless competition dynamics. A: Variance of L23E firing rate within trials for different combinations of local self-excitation (*J_local_*) and lateral-inhibition strengths (*J_lateral_*) without (top) and with (bottom) stimulus drive in L4. Variance of the firing rate reveals the winnerless competition regime (WL) because firing rates in the non-competing (NC) and winner-take-all (WTA) regimes do not change over time while firing rates in the WL regime alternate between high and low rates. Local self-excitation and lateral-inhibition in L23 both facilitate competition. The model can occupy 3 states: non-competing (NC), winnerless dynamics (WL) and winner-take-all (WTA). Without feedforward stimulation in L4 the model does not compete (NC) (top). Applying an input of 20 Hz reveals a bistable regime where the variance increases (WL; bottom). The red dot indicates the parameter settings used in further simulations. B: Dominance duration for different combinations of feedforward input in L4 to the two columns. right: Dominance duration per column when changing input to one column while keeping input to the other column constant. C: Alternation rate for different combinations of feedforward input in L4 to the two columns. right: Increasing total input decreases alternation rate. Alternation is highest when columns are equally stimulated.

We expect the model to align well with all four of Levelt’s propositions. We first test whether the first three propositions are reproduced by the winnerless competition resulting from applying stimulation to L4 of both columns. Specifically, we expect the dominance duration to primarily increase for the column receiving stronger stimulation. Furthermore, we expect the alternation rate to be maximal when both columns are equally stimulated. Each column receives an external input applied through 300 and 188 synapses to the excitatory and inhibitory populations of L4, respectively (ratio derived from [47]), resulting in a net excitatory stimulation. We simulate 10 trials for each condition with 100 seconds per simulation. From each simulation we extract the dominance duration per column and the alternation rate. The winner is the column whose excitatory L23 population exhibits the highest firing rate. We define a dominance duration interval as the time between two successive switches of dominance. Increasing the stimulus drive to one column mainly increases the dominance duration of that column (Fig 3B). In line with our hypotheses, the alternation rate is maximal when stimulation of the columns is equal and decreases when the difference in stimulation is increased (Fig 3C). The alternation rate is decreased with stronger total stimulus drive, compared to the behaviorally observed increase. The alignment of the model with the first three propositions shows that feedforward stimulation of L4 affects the mutual-inhibition mechanism in L23 by increasing the depth of the attractor receiving the highest stimulation. Without input to deep layers, the model does not account for the fourth proposition. Indeed, as was found for other winnerless competition models, the alternation rate is not enhanced with higher stimulus drive because it increases competition between L23 populations and deepens the attractors, lowering the probability of transitions. Up to now we only applied feedforward stimulation to L4. However, applying no stimulation to deep layers is not biologically realistic [2, 8, 17, 27, 28, 52]. Therefore, in the next section we discuss the influence of deep layers on winnerless competition dynamics.

### External Input to Deep Layers explains Levelt’s Fourth Proposition

We expect that deep layers 5 and 6 implement a gain control mechanism on the upper layers [10, 21, 43, 44]. Such a gain control mechanism would promote shallow attractors and enhance the alternation rate with increased stimulus drive. We apply external input to L5 and L6 through all external synapses (see 1) to excitatory and inhibitory populations, (Fig 4A), resulting in a net excitatory input. We systematically vary the stimulus drive to deep layers L5 and L6 and obtain the alternation rate and dominance duration are for each combination. Externally stimulating L5 or L6 both increases the alternation rate (Fig 4B) and decreases the dominance duration (Fig 4C) of the winnerless column model. Externally stimulating L5 and L6 simultaneously further enhances this effect. These results indicate that layer-specific stimulation of deep layers during bistable perception result in output matching Levelt’s fourth proposition: increased alternation rate in response to increased stimulation. Effects are generally larger for stimulation of L5, but robust for stimulation of L6 as well. External input can target either or both excitatory and inhibitory populations within a layer. To show how the excitation-inhibition balance of the layer-specific input affects the alternation rate we systematically apply different levels of input to the excitatory and inhibitory populations of the layers in question, L5 (Fig 4D) and L6 (Fig 4E). For both layers we find that more excitation increases higher alternation rate. More inhibitory input would not affect the alternation rate significantly. Interestingly, for L6 there is a slight decrease before the alternation rate starts increasing when the layer is excited more. Together these results suggest that deep layers modulate the winnerless dynamics. Stimulation of excitatory populations in these layers causes an increase in alternation rate which aligns the winnerless dynamics of the model with all of Levelt’s propositions for bistable perception, including the fourth.

**Fig 4.**
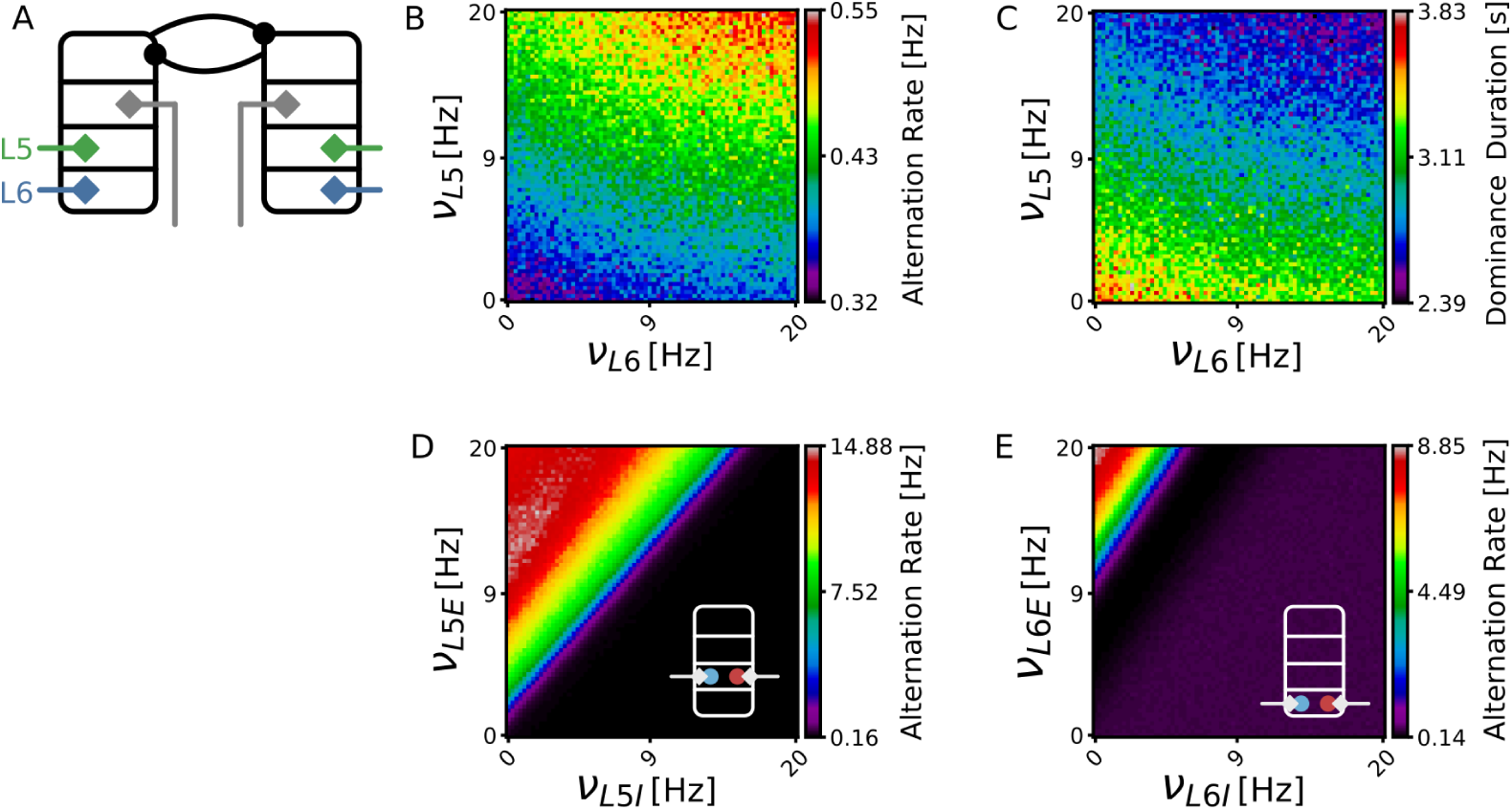
External input to deep layers. A: External input is applied to deep layers L5 and L6. Combinations of input to L5 and L6 increase alternation rate (B) and reduce dominance duration (C). External input to excitatory or inhibitory populations in L5 (D) and L6 (E). More excitatory input mainly increases the alternation rate in both layers.

### External Input to Granular Layer explains Levelt’s Fourth Proposition

Several models of bistable perception suggest a form of input suppression to account for Levelt’s fourth proposition [15, 18, 29]. Such architectures generally consider a two-level hierarchy where the input components are isolated from the decision making components. The decision making components then inhibit the input components and thereby sustain shallow attractors. The laminar structure of our model naturally provides such a split structure. Granular layer L4 generally receives stimulus input while L23 implements a decision making mechanism through the lateral mutual-inhibition mechanism between two columns. To test whether the interlaminar connectivity between L4 and L23 implements an input suppression mechanism we examine various combinations of external stimulation of L23 and L4 and their effect on the alternation rate and dominance duration. We apply external input to L23 and L4 through all external synapses (see 1) targeting excitatory and inhibitory populations (Fig 5A), resulting in a net excitatory input. We systematically vary the external input to superficial L23 and granular L4 used in simulations and record the corresponding alternation rates and dominance durations. Injecting external input to the decision layer L23 generally decreases the alternation rate (Fig 5B) and increases the dominance duration (Fig 5C). This reflects a deepening of the attractors, which is common in simple mutual-inhibition network models of bistable perception which consists of only a decision layer [12]. For external input to the granular layer L4 the results are less straightforward. For weak inputs to L4 the alternation rate decreases (Fig 5B) and dominance duration increases (Fig 5C), just as it does for inputs to L23. Interestingly, increasing external input to L4 further reverses the effect and causes the alternation rate to increase again. This non-linear relation between the external input and the alternation rate reflects first a deepening and than a diminishing of the attractor for increasing stimulus drive. This suggests a suppression of L4 activity when external input to L4 is strong. To explore how the excitation-inhibition ratio of the layer-specific input affects the alternation rate we perform a grid search applying different levels of input to the excitatory and inhibitory populations of the layers in question, L23 (Fig 5D) and L4 (Fig 5E). We find that more inhibitory input to superficial layer L23 increases the alternation rate, which is expected as inhibitory input would diminish the competition between columns by reducing the attractor depth. For granular layer L4 we find that it is particularly unbalanced input (i.e., more excitatory or more inhibitory) which increases the alternation rate while for balanced input (i.e., similar excitatory as inhibitory input) the alternation rate is at its lowest point. This confirms again that more excitatory input diminishes the depth of the attractors, but in addition shows that predominantly inhibitory input to L4 has a similar effect. Feedforward stimulation between areas is generally of excitatory nature and generally targets L4. Our results suggest that such unbalanced excitatory input could very well cause an increase in the alternation rate.

**Fig 5.**
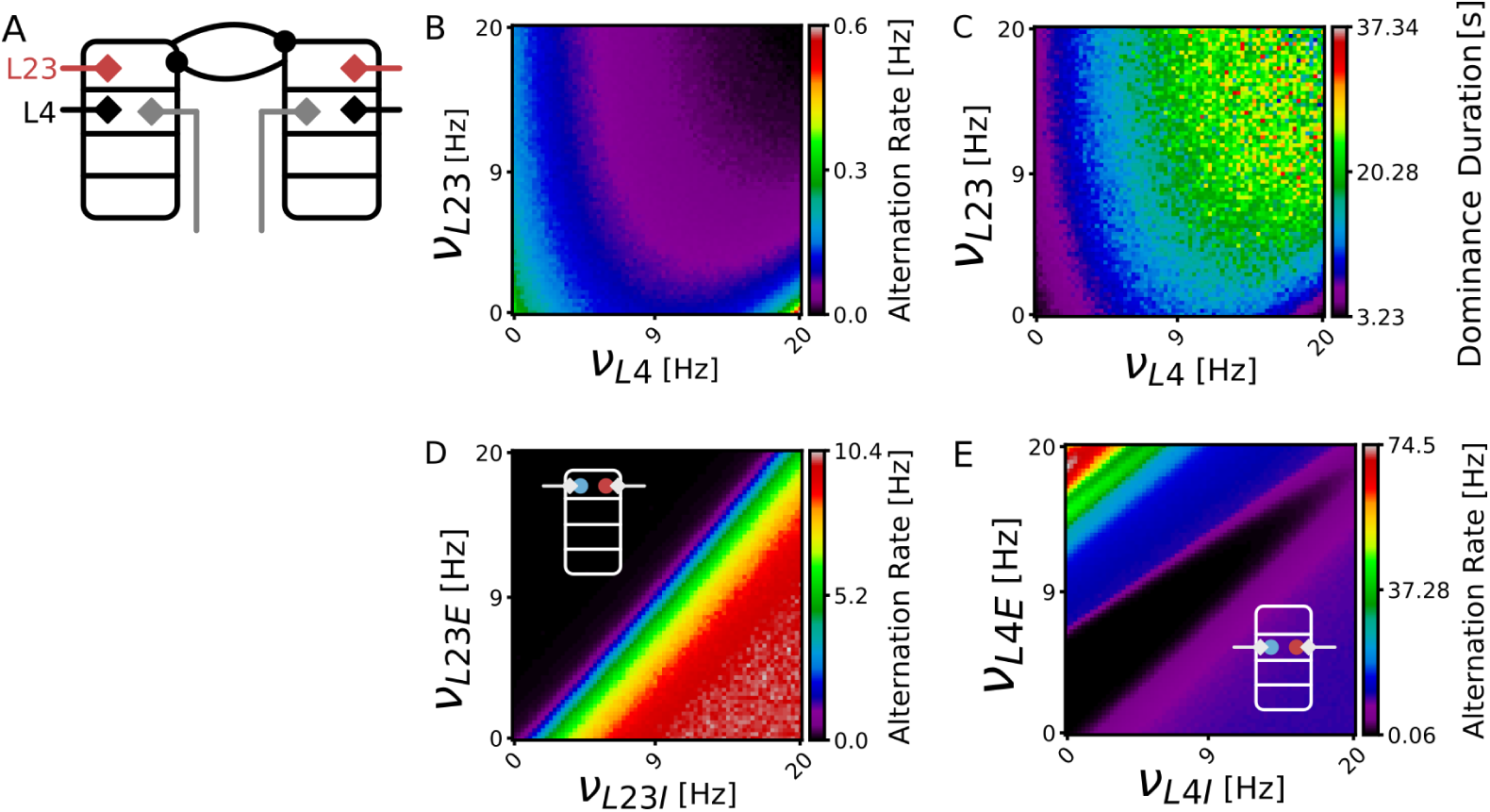
External input to superficial and granular layers. A: External input is applied to layers L23 and L4. B: Alternation rate decreases with increasing excitatory input to L23. For relatively strong L23 input, alternation rate is decreased for weak input to L4 as well. C: Dominance duration increases with increased excitatory L23 input. Dominance duration increases for weak input to L4 but decreased for strong stimulation of L4. D: External input to excitatory or inhibitory populations in L23. More inhibitory input mainly increases the alternation rate. E) External input to excitatory or inhibitory populations in L4. More unbalanced input increases the alternation rate. The effect is more pronounced for strong excitatory input.

### Interlaminar Mechanisms underlying Levelt’s Fourth Proposition

Our results from simulations wherein we apply varying amounts of external input to different layers support the hypotheses that deep layers implement gain control while superficial and granular layers conjointly implement an input suppression mechanism. We argue that these mechanisms critically depends on the interlaminar connectivity profile we derived from biological datasets. Here, we explore this further and elucidate the specific interlaminar circuits of the columnar model that contribute to gain control and input suppression.

#### Gain Control by Deep Layers

Potjans & Diesmann [47] identified a feedforward flow of activity resulting from the interlaminar connectivity of the circuit: L4 excites L23, which in turn excites L5. Layer 5 then excites L6. Finally, L5 and L6 inhibit L23 and L4, respectively. Deep layers balance the circuit by directly targeting the superficial and granular layers with interlaminar connections. We explore whether these connections are also directly responsible for modulations in alternation rate and dominance duration by manipulating their strength. We control the connection strength of all connections from deep layer populations L5 and L6 onto superficial and granular layers, L23 and L4 respectively (Fig 6A). Completely uncoupling the deep layers from the upper layers decreases the alternation rate (Fig 6A). That is, without the inhibiting effects of the deep layers, the dominant attractor state becomes deeper, resulting in increased dominance duration and less alternations between states (Fig 6A,B). Increasing the connection strength from deep layers to upper layers increases the alternation rate and suppresses the dominance duration in a linear fashion. Together, these results show that it is the direct interlaminar connections of deep layers onto upper layers L23 and L4 which affect the winnerless dynamics and give rise to statistical regularities summarized in Levelt’s Fourth proposition. Consistent with empirical findings [10, 21, 43, 44], the deep layers within this columnar model exhibit an inhibitory effect on the upper layers. We establish a direct connection between these biological findings and behavioral observations of statistics related to bistable perception.

**Fig 6.**
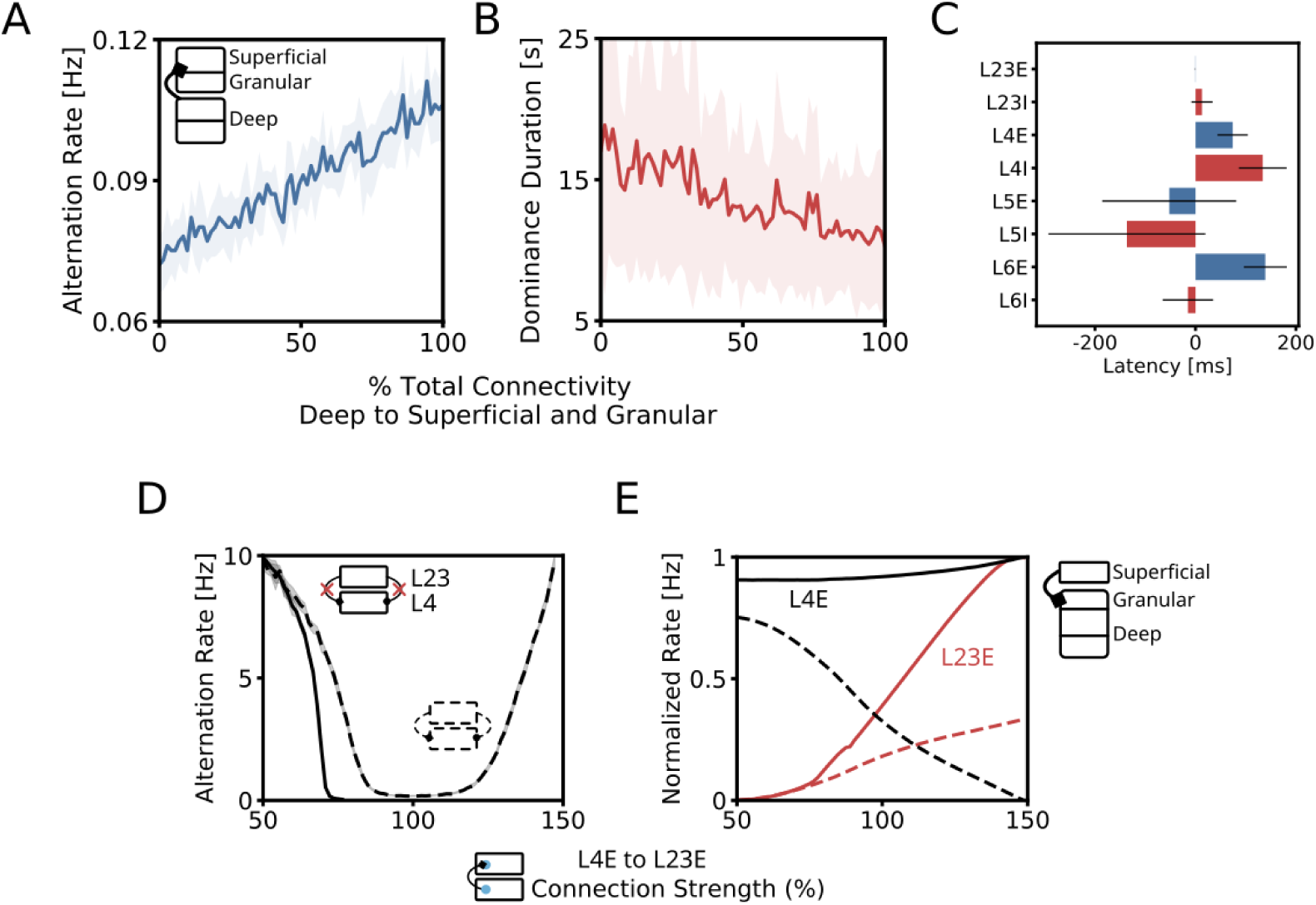
Exploring interlaminar mechanisms. A: Alternation rate is increased by increasing connection strength of deep layers to upper layers. B: Dominance duration is decreased by increasing connection strength of deep to upper layers. C: Population specific latency relative of switching attractor states in L23E. Lower latency indicates a early detection of switching states compared to population L23E. Latency measured for all decision reversals for 100*s* simulations for 10 trials. Black horizontal bars indicate standard deviation. D: Effect of L23 to L4 connections on the alternation rate in a superficial-granular sub-circuit. Increasing weak feedforward connectivity (L4E to L23E, 50 *−* 100%) decreases the alternation rate in both the circuit with L23 to L4 connections (dashed) and the circuit without these connections (solid). Increasing stronger feedforward connectivity (100 *−* 150%) the alternation rate of the circuit with L23 to L4 connectivity increases while the alternation rate of the other circuit does not. E: Effects of L23 to L4 connections on the mean rate of L23E (red) and L4E (black) populations. Rates are normalized per population.

To further investigate the causal role of deep layers in decision reversals we compare the timing of state switching in the deep and superficial layers. Specifically, for each decision reversal from one attractor state to the other we take a 400*ms* interval around the time-point where the reversal took place in L23E (i.e., when the rate of L23E in the dominant column first dropped below the rate of L23E in the other column). The population rates in this interval are then used to obtain the population specific reversal point (i.e., when the rate of a particular population in the dominant column first dropped below the rate of the same population in the other column). Interestingly, L5 populations show a particularly early onset of switching decision compared to L23 (Fig 6C). This suggests a causal influence of L5, and in particular the inhibitory population, in bringing about decision reversals. An early review on the laminar and lateral structure of cortical circuits assigns a central role to L5 in decision making [19]. According to the authors, L5 would confirm the decision *after* it was made by superficial layers and prepare the cortical output. However, our results suggests that L5 might causally affect the decision process in the superficial layers and play an important role in reversing rather than confirming a decision. We suggest that L5 ensures that attractor states remain shallow, thereby promoting exploration of the attractor landscape. The switches in L6 excitatory population are relatively late compared to decision layer L23. L6 inhibitory neurons switch at similar times as L23. We suggest these results on the population-specific latency of switches only provide information on whether a population is involved in causing particular jumps across the attractor separatrix and are independent from the effects of constant input, as this mainly affects the attractor depth during the simulation.

#### Input Suppression in Superficial-Granular Circuits

We find that weak input to granular layer L4 deepens the attractors while strong input renders them more shallow, promoting reversals between states. We hypothesize that L23 and L4 implement an input suppression mechanism where L23 suppresses L4 [14]. To inspect whether interlaminar connections from L23 onto L4 are indeed capable of causing the observed increase in alternation rate for strong input to L4, we first uncouple the deep layers (L5 and L6) from the superficial and granular layers. Next, we compare two implementations of this superficial-granular sub-circuit: One with and without L23 connections targeting L4. We then systematically manipulate the feedforward connection from L4E to L23E which controls the feedforward input to the decision layer L23. We keep the external input fixed. For weak feedforward connectivity (*<* 100%) the alternation rate strongly decreases for both versions of the circuit, indicating a deepening of the attractors (Fig 6D). However, for strong feedforward connections (*>* 100%) the circuit implementing direct feedback connections from L23 onto L4, the alternation increases. This suggest connections from L23 onto L4 ensure shallow attractors and promote reversals. To show that L23 is suppressing its own input, we investigated the effect that the feedback connections from L23 to L4 have on the mean firing rate of these populations. Specifically, we implement two versions of a full column model. One with and one without connections from L23 to L4. As before, we manipulate the feedforward connection strength, L4E to L23E, while keeping the external input fixed. We compare the normalized rates of L23E and L4E in both versions of the column. Without connections from L23 to L4 it is mainly the rate of L23E which is increased while the rate L4E is not much affected (Fig 6E). With L23 to L4 connections the increase in rate of L23E is much weaker. Interestingly, the rate of L4E is now strongly affected by an increase in feedforward connectivity. The L23 to L4 connections cause a strong decrease in L4E rate. These results show that L23 and L4 in the cortical column model implement an input suppression mechanism through direct feedback connections from L23 to L4 within the same column.

## Discussion

We test the significance of the layered structure of the cortex for bistable perception. Using a dynamic mean field model of two cortical columns with empirically derived interlaminar circuit connectivity, we show that the architecture of the cortical microcircuit provides the neurobiological substrate required for winnerless competition dynamics supplemented with gain control and input suppression mechanisms. Two columns in our model compete for dominance through mutual-inhibition in superficial layers L23. We show that the first three of Levelt’s propositions are directly accounted for by the model through systematically applying feedforward stimulation to the main input layer L4 of both columns. Furthermore, we establish that concomitant input to deep layers increases the alternation rate as a function of stimulus drive. This shows that feedforward input to deep layers accounts for Levelt’s fourth proposition using a mechanism grounded in biology. In line with theoretical work on bistable perception [40], we suggest that deep layers implement a gain modulation mechanism on the upper layers through interlaminar inhibition. This requires that deep layers receive stimulus-related feedforward input, which is supported by several studies that have shown that deep layers are important targets for thalamic axons (both matrix and core thalamic nuclei) in many brain areas [2, 8, 17, 28, 52]. Our model further shows that relatively weak inhibition of the upper layers is already sufficient to increase the alternation rate, however several studies have highlighted that inhibitory effects of layer 6 on the upper layers are rather strong [10, 21, 43, 44]. These studies are based on mouse cortex while our model implements connectivity derived mainly from cat. Furthermore, [10, 21] have shown that the mechanism responsible for such gain modulation involves a more complex inhibitory circuit than currently implemented in our *Laminar Column Model*. The wide variety of cell types and projections suggest a variety of layer 6 dependent circuits with different roles. Further research is needed to delineate how layer 6 is involved in the implementation of gain control [43]. We suggest that extending our model to include a more sophisticated inhibitory circuit in layer 6 would amplify the importance of this layer in modulating the winnerless competition dynamics and in maintaining shallow attractors. Nevertheless, our model is capable of implementing gain control due to weak inhibitory connections from deep layers to superficial and granular layers. Because empirical and modelling work suggests that deep layers inhibit upper layers we hypothesize that projections from deep to upper layers are responsible for maintaining shallow attractors required for high alternation rates. By systematically modulating the connectivity from deep to upper layers we show that it is indeed these connections that are responsible for increasing the alternation rate.

Other theories of bistable perception have postulated forms of input suppression to explain Levelt’s fourth proposition [15, 18, 25, 29]. We hypothesize that local recurrent feedback from L23 to L4 implements such a mechanism. We find that applying weak input to L4 (*<* 9*Hz* to L4 and no input to L23) reduces the alternation rate, but when stronger input is applied (*>* 9*Hz*) the alternation rate starts to increase again. By systematically modulating the recurrent feedforward and feedback connections between L23 and L4 we show that it is indeed the feedback from L23 onto L4 which causes this increase in alternation rate during strong stimulation of L4. With this we show that the columnar circuit implements a form of input suppression that can account for the increase in alternation rate when the stimulation is strong. Interestingly, one study has shown a potential deviation from Levelt’s fourth proposition as formulated in [11]. By alternating the contrast and motion strength separately in a random dot motion experiment [46], these authors found a decrease of alternation rate for weak motion strength (but not for weak contrast levels) and an increase in alternation rate for strong motion. Importantly, unlike previous models, our model accounts for this deviation from Levelt’s fourth proposition at weak stimulus drives. We show that weak feedforward stimulation to L4 reduces the alternation rate and strong stimulation increases the alternation rate matching the experimental observation in [46].

Our findings have important implications for a systems-level perspective on bistable perception as well. The layer-specificity of the gain control and input suppression mechanisms in conjunction with characteristic laminar patterns of feedforward, lateral and feedback projections in the cortex, allows us to draw inferences and make predictions on how cross-regional interactions affect bistable perception. For example, top-down feedback selectively targets deep layers during viewing of ambiguous figures [32]. Furthermore, activity in frontal areas is associated with higher alternation rates [60]. Together with insights gleaned from our model, these observations support the hypothesis that increased top-down feedback from higher cortical areas targeting deep layers in visual cortex serves to inhibit the upper layers in order to maintain shallow attractors and increase the alternation rate. Through such a mechanism the frontal areas, known to be important for executive control and voluntary action [22, 39], may actively induce switches. Furthermore, many theoretical studies highlight the importance of top-down and bottom-up interactions for bistable perception without specifying the neural substrate that would implement these mechanisms [15, 18, 25, 29]. We show that interlaminar circuit implements mechanisms which explains all of Levelt’s propositions and thus provides a possible neural substrate for bistable perception locally within a brain region. How much local and inter-area mechanisms each contribute to bistable perception remains an open question, but our work highlights the importance of the local inter-laminar circuitry. Further extensions of our model to encompass additional brain regions are warranted to gain a more comprehensive understanding of the interplay between local and inter-area inhibitory effects. Future work on layer specific responses and connectivity during bistable vision could help to further constrain our model. For example, the advent of ultra-high field fMRI allows us to investigate neural correlates of human cognition at columnar and laminar resolution [30, 45].

In conclusion, previous work has provided important insights how bistable perception may possibly be implemented algorithmically [15, 18, 25, 29, 40]. Our work provides a first crucial step towards an understanding which and how these algorithms are implemented by the neurobiological hardware of the cortex [31, 37]. Our study provides compelling evidence for the relevance of gain modulation and input suppression mechanisms for bistable perception and the implementation of these mechanisms in distinct laminar circuits.

## Methods

### Dynamic Mean Field

We use a dynamic mean field (DMF) implementation which simulates neural populations of the cortical column. A single column consists of 8 populations divided across 4 layers: L23, L4, L5 and L6 1. Each layer contains an excitatory and an inhibitory population. The intra- and interlaminar connectivity is derived from [47] which base their connectivity scheme on empirical research reviewed in [8, 55]. We adjust the population sizes to fit with area MT of macaque using estimated population sizes computed by [50]. The empirical nature of our model necessitates that we implement a column of a specific cortical region. We chose MT because a recent study showed that two mutually exclusive interpretations of an ambiguous motion stimulus were reflected by distinct cortical columns in (human) MT. However, the mechanism we identify generalize to other cortical regions (see Supplementary Materials for layer specific input using connectivity of other visual system areas). We implement two columns, each representing an axis-of motion column in MT, one for each direction of motion (horizontal (H) or vertical (V)). The firing rate dynamics of each population are modelled using a set of differential equations. Each population has five state variables: the synaptic current, the membrane potential, noise, an adaptation variable and the firing rate. The change in current 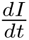 to each population in the DMF model is given by the stochastic differential equation:

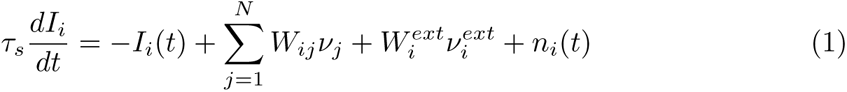

Where *τ_s_*is the synaptic time constant, *W* and *ν* the recurrent connectivity strength and the recurrent input from other populations, respectively. *W^ext^*and *v^ext^*are the external weight and input to population *i*. Finally, the noise *n*(*t*) is an Ornstein-Uhlenbeck process [23] with zero mean and deviation *σ*:

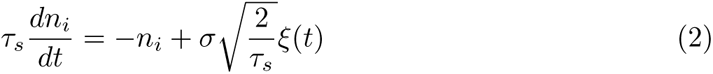

Where *ξ*(*t*) is a white noise process with zero mean and variance 1. The current is passed through a linear temporal filter to obtain the change in population membrane potential:

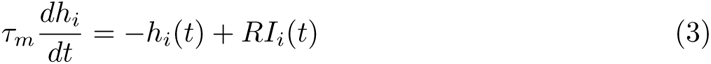

Where *τ_m_* is the membrane time constant and *R* is the membrane resistance given by 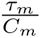. Adaptation is modelled by updating an rate-dependent variable (*w*) with a slow time-constant:

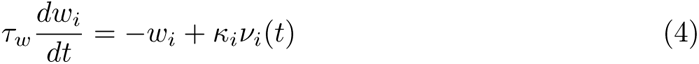

Where *τ_w_*, *κ_i_* and *ν_i_*(*t*) are the adaptation time constant and population specific adaptation strength and firing rate, respectively. Finally, the membrane potential and the adaptation variable are passed through a non-linear threshold function [61] to obtain the population firing rate:

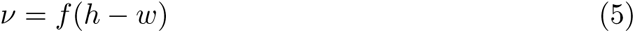

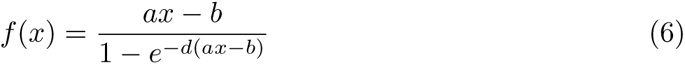

Where *a*, *b* and *d* are the gain, threshold and noise factor, respectively. These parameters are fitted such that the DMF and LIF implementation of the microcircuit model have similar firing rates (see 1). Code for simulating the dynamic mean field is available at github.com/ccnmaastricht/LCM-BS

### Network connectivity

We obtain the intra- and interlaminar recurrent connection probabilities from [47]. The feedforward connection probability from L4E to L23E is doubled as in to compensate for the decrease in firing rate in L23E due to the lateral inhibition (c.f. [14, 59]). We subsequently compute the number of synapses between populations:

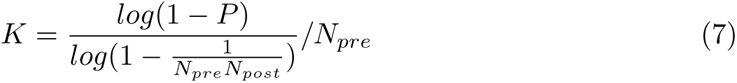

Where *P*, *N_pre_* and *N_post_* are the connection probabilities between the pre- and post-synaptic populations and the number of neurons in the pre- and post-synaptic populations, respectively. The layer-specific excitation-inhibition ratio is obtained by dividing the number of excitatory neurons by the inhibitory neurons. The inhibitory synaptic weight is then the multiplication of this ratio with *−J_E_*. The final connectivity matrix is obtained by multiplying the number of synapses by the synaptic weight:

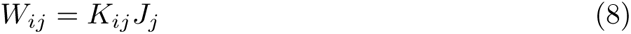

External input is applied by defining a number of synapses and an input frequency and multiplying the these with the standard excitatory weight *J_E_*. Each population in the network receives a background input with a frequency of 8 Hz via a population specific number of synapses with weight *J_E_* = 0.0878. The number of background synapses are taken from [47]. Lateral mutual-inhibition is modelled by symmetric connections from population L23E in one column to L23I in the other column. *J_local_*and *J_lateral_*control the self-excitation of L23E and lateral-inhibition strength, respectively. Feedforward stimulation of L4 excitatory and inhibitory populations is applied through 295 and 186 synapses, respectively (with the same E/I-ratio as [47]). The number of external connections of net excitatory external inputs in layers L4, L6 and layers L23, L5 are 300 and 255 for excitatory and inhibitory populations, respectively. The full list of model parameters are provided in 1 and 2. Fig 2 shows a diagram of the implemented model.

**Table 1.** Model Structure.

|  |  |  |  |  |  |  |  |  |
| --- | --- | --- | --- | --- | --- | --- | --- | --- |
| Populations | 16 populations, 2 columns, 8 per columns |  |  |  |  |  |  |  |
| Num. Synapses | $K$, within, between columns, stimulus drive | | | | | | | |
| Weights | $J$, within and between columns | | | | | | | |
| Population size | $N$, area and population specific | | | | | | | |
|  | L23E | L23I | L4E | L4I | L5E | L5I | L6E | L6I |
| Population Sizes |  |  |  |  |  |  |  |  |
| $N_{V1}$ | 20683 | 5834 | 21915 | 5479 | 4850 | 1065 | 14395 | 2948 |
| $N_{MT}$ | 30303 | 8547 | 14101 | 3525 | 7088 | 1556 | 7918 | 1621 |
| Number of External Synapses |  |  |  |  |  |  |  |  |
| $K_{bg}$ | 1600 | 1500 | 2100 | 1900 | 2000 | 1900 | 2900 | 2100 |
| $K_{ff}$ | 0 | 0 | 295 | 186 | 0 | 0 | 0 | 0 |
| $K_{ext}$ | 300 | 255 | 300 | 255 | 300 | 255 | 300 | 255 |
| $\kappa$ | 1.5 | 0.0 | 0.0 | 0.0 | 0.0 | 0.0 | 0.0 | 0.0 |
| Columnar Connectivity |  |  |  |  |  |  |  |  |
| $P_{column}$ | source: | | | | | | | |
| target: | L23E | L23I | L4E | L4I | L5E | L5I | L6E | L6I |
| L23E | 0.1009 | 0.1689 | 0.0880 | 0.0818 | 0.0323 | 0.0000 | 0.0076 | 0.0000 |
| L23I | 0.1346 | 0.1371 | 0.0316 | 0.0515 | 0.0755 | 0.0000 | 0.0042 | 0.0000 |
| L4E | 0.0077 | 0.0059 | 0.0497 | 0.1350 | 0.0067 | 0.0003 | 0.0453 | 0.0000 |
| L4I | 0.0691 | 0.0029 | 0.0794 | 0.1597 | 0.0033 | 0.0000 | 0.1057 | 0.0000 |
| L5E | 0.1004 | 0.0622 | 0.0505 | 0.0057 | 0.0831 | 0.3726 | 0.0204 | 0.0000 |
| L5I | 0.0548 | 0.0269 | 0.0257 | 0.0022 | 0.0600 | 0.3158 | 0.0086 | 0.0000 |
| L6E | 0.0156 | 0.0066 | 0.0211 | 0.0166 | 0.0572 | 0.0197 | 0.0396 | 0.2252 |
| L6I | 0.0364 | 0.0010 | 0.0034 | 0.0005 | 0.0277 | 0.0080 | 0.0658 | 0.1443 |
| $J_{column}$ | 0.0878 | -.3113 | 0.0878 | -.3512 | 0.0878 | -.3998 | 0.0878 | -.4287 |
| Lateral Inhibition Parameters |  |  |  |  |  |  |  |  |
| $P_{lateral}$ | 0.1 | | | | | | | |
| $J_{lateral}$ | 0.172 | | | | | | | |
| $J_{local}$ | 0.13 | | | | | | | |
| $P$: connection probabilities; $J$: synaptic strength (mV); $\kappa$: adaptation strength | | | | | | | | |

### 0.1 Simulation

All simulations of the stochastic differential equations describing our model are performed in Python 3.8.10 using the Euler-Maruyama method with a simulation time of 1.0 s and a time step of 0.1 ms. Simulations are executed on the PizDaint CSCS Supercomputer cluster.

**Table 2.** Model Parameters.

| Dynamic Mean Field Parameters |  |  |
| --- | --- | --- |
| Name | Value | Description |
| $\sigma$ | 0.02 nA | noise amplitude |
| $\tau_s$ | 0.5 ms | synaptic time constant |
| $\tau_m$ | 20.0 ms | membrane time constant |
| $C_m$ | 250.0 mF | membrane capacitance |
| $a_f$ | 48 | function gain |
| $b_f$ | 981 | function threshold |
| $d_f$ | 0.0089 | function noise factor |
| $\nu_{bg}$ | 8 Hz | background rate |
| $\tau_w$ | 10 s | adaptation time constant |

### 0.2 Analysis

Our main analyses focus on the distribution of the dominance duration and alternation rate. The dominance duration is defined as the length of the time intervals the model occupies one of the attractors (i.e. L23E rate of column D1 is high and D2 is low, or vice versa). To obtain the distribution of dominance durations, we extract the reversal time-points of L23E populations by comparing their rates. The difference between consecutive reversal time-points quantifies the dominance duration.

The alternation rate is here defined as the number of reversals per second per simulation (in Hz). Thus, every instance of a simulation has a single value for the alternation rate. Distributions for the alternation rate are obtained by running multiple trials using different random noise realizations.

## Acknowledgments

This study has received funding from the European Union’s Horizon 2020 Framework Programme for Research and Innovation under the Specific Grant Agreement No. 945539 (Human Brain Project SGA3). We would like to thank Alessandra Pizzuti the helpful discussions on bistable perception and ultra-high field fMRI. Finally, we acknowledge the use of Fenix Infrastructure resources, which are partially funded from the European Union’s Horizon 2020 research and innovation programme through the ICEI project under the grant agreement No. 800858.

## Author Contributions

**Conceptualization:** Kris Evers, Mario Senden. **Investigation:** Kris Evers. **Methodology:** Kris Evers, Mario Senden. **Project Administration:** Kris Evers, Judith Peters, Rainer Goebel, Mario Senden. **Software:** Kris Evers. **Supervision:** Mario Senden, Judith Peters. **Visualization:** Kris Evers. **Funding Acquisition:** Rainer Goebel **Writing - Original Preparation:** Kris Evers. **Writing - Review & Editing:** Kris Evers, Judith Peters, Mario Senden.

