## Supplementary figures and images for "Layered Structure of Cortex Explains Reversal Dynamics in Bistable Perception"

### Supplemental Figure 1

Supplementary Material

S1 Fig. Layer Specific External Input in Areas of the Visual System

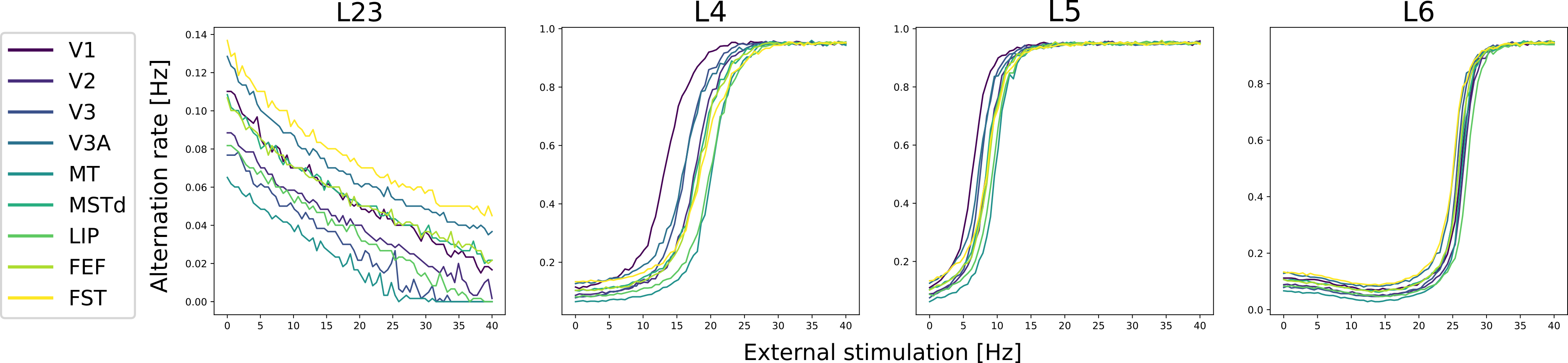
